# ‘Mind *in Vitro*’ platforms: Versatile, scalable, robust and open solutions to interfacing with living neurons

**DOI:** 10.1101/2023.08.21.554033

**Authors:** Xiaotian Zhang, Zhi Dou, Seung-Hyun Kim, Gaurav Upadhyay, Daniel Havert, Sehong Kang, Kimia Kazemi, Kai-Yu Huang, Onur Aydin, Raymond Huang, Saeedur Rahman, Austin Ellis-Mohr, Hayden A. Noblet, Ki H. Lim, Hee Jung Chung, Howard J. Gritton, M. Taher A. Saif, Hyun Joon Kong, John M. Beggs, Mattia Gazzola

**Affiliations:** Carl R. Woese Institute for Genomic Biology, University of Illinois at Urbana–Champaign, Urbana, IL 61801; Department of Mechanical Science and Engineering, University of Illinois at Urbana–Champaign, Urbana, IL 61801; Department of Physics, Indiana University Bloomington Bloomington, IN 47405; Department of Chemical and Biomolecular Engineering, University of Illinois at Urbana–Champaign, Urbana, IL 61801; Department of Electrical and Computer Engineering, University of Illinois at Urbana–Champaign, Urbana, IL 61801; Molecular and Integrative Physiology, University of Illinois at Urbana–Champaign, Urbana, IL 61801; Neuroscience Program, University of Illinois at Urbana–Champaign, Urbana, IL 61801; Beckman Institute for Advanced Science and Technology, University of Illinois at Urbana–Champaign, Urbana, IL 61801; Department of Comparative Biosciences, University of Illinois at Urbana–Champaign, Urbana, IL 61802

## Abstract

Motivated by the unexplored potential of *in vitro* neural systems for computing, and by the corresponding need of versatile, scalable interfaces for multimodal interaction, we present an accurate, modular, fully customizable and portable recording/stimulation solution that can be easily fabricated, robustly operated, and broadly disseminated. A reconfigurable platform that works across multiple industry standards enables a complete signal chain, from neural substrates sampled through high-density Micro-Electrode Arrays (MEAs) to data acquisition, downstream analysis and cloud storage. Built-in modularity supports the seamless integration of electrical/optical stimulation and fluidic interfaces. Custom MEA fabrication leverages maskless photolithography, favoring the rapid prototyping of a variety of configurations and spatial topologies. Through a native analysis and management software suite, the utility and robustness of our system is demonstrated across neural cultures and applications, including embryonic stem cell-derived and primary neurons, organotypic brain slices, 3D engineered tissue mimics, concurrent calcium imaging and long-term recording. Overall, our technology, termed ‘Mind *in Vitro*’ to underscore the computing inspiration, provides an end-to-end solution that can be widely deployed due to its affordable (>10X cost-reduction) and open-source nature, catering to the expanding needs of both conventional and unconventional electrophysiology.

## 1 Introduction

Neural tissue supports a host of information processes fundamental to many organisms, from autonomic body functions to motion, sensing and high-level reasoning [1, 2, 3]. In the quest to decode the inner workings of neural architectures, their inspection has long relied on electrical recordings [4, 5]. While *in vivo* electrophysiology, instrumental in neuroscience, uniquely allows for sampling neural activities associated with specific behaviors [6, 7], its interpretation is challenged by whole-organism complexity. *In vitro* systems, including single cells, small networks, or tissue samples of larger-scale connectivity, in contrast represent a reduced –yet complementary– route to expose neural interactions across scales [8, 9, 10, 11, 12, 13]. From the perspective of synthesizing fundamental principles of biological computing, there are significant opportunities with deploying electrical interfaces *in vitro*, particularly in conjunction with engineered neural substrates [11, 12, 13, 14]. Indeed, by spatially distributing and connecting biological units (neural populations) of prescribed size, geometry or neuron-type onto input/output electronic platforms, living processing architectures may be realized, operated and tested [15, 16, 17, 18, 19], potentially enabling a new class of computing systems, with ramifications in engineering, biology and healthcare.

In this context, Micro-Electrode Arrays (MEAs) technology, where electrodes of variable size, density and spatial arrangement are patterned on biocompatible surfaces, represents a powerful and mature option to interface with cellular systems, and has been employed from single neurons electrophysiology [20] to high-throughput pharmaceutical screens [21, 22]. However, the potential of MEA systems remains hindered by design, fabrication, integration and software management complexities. Labs have indeed little choice but to invest in commercial solutions [23, 24], which are proprietary, specialize to support biomedical research, afford minimal-to-no customization, and remain costly, limiting adoption (from tens of thousands of dollars for a standard 60-electrode system to hundreds of thousands for more advanced configurations). This has led to a recent interest in the open-source development of MEA platforms [25, 26], aimed at lowering barriers to entry while catering to needs such as multi-well drug testing [27], enhanced optical access [28, 29], or chronic tissue monitoring [26]. However, a comprehensive, accessible solution to flexibly and multimodally interact with a variety of neural substrates, from organotypic brain slices to engineered 3D tissue, remains to be demonstrated.

Here, prompted by the untapped potential of *in vitro* neural systems for computing, we present an interfacing platform that is versatile, scalable, reconfigurable and portable, that can be easily fabricated, robustly operated across cellular contexts, and broadly disseminated. We term this technology ‘Mind *in Vitro*’ (MiV), to emphasize the information processing motivation. Our platform hosts high-density MEA chips, manufactured in standard cleanroom facilities via maskless photolithography, and varying in size, spatial topology and transparency depending on the application. These chips are matched to swappable, custom printed circuit boards (PCBs) relaying neural signals to Open Ephys [30] or Intan acquisition terminals [31] for signal acquisition and subsequent downstream processing. Our system seamlessly integrates with both electrical and optical stimulation modules, as well as other add-ons such as fluidic interfaces or tissue-specific positioning apparatuses. Reconfigurability is further leveraged to comply with industrial standards and integrate with common microscopic chambers, enabling concurrent imaging. Such flexibility allows for the combination of high-temporal (electric) and high-spatial (imaging) resolutions, seizing on the opportunities afforded by ever-expanding genetic optical markers used in neuroscience research [32, 33]. Additionally, an open-source software package is developed to manage the system and support operability, data storage, analysis, and visualization. To increase usability, source code is based on Python language and native interfaces with external neurophysiology (NeuralEnsemble [34]) and machine learning (scikit-learn [35]) software suites are provided.

Our integrated systems are demonstrated across a broad range of *in vitro* settings, from 2D cultures of embryonic stem cell-derived neurons and dissociated hippocampal cells to organotypic brain slices and 3D engineered neural tissue mimics. Multiple applications are illustrated including electrical, optical and fluidic manipulation, concurrent calcium imaging, and long-term recording (*>*24h). By logging over 1000 hours of experiments and tens of TB of data across distinct labs, robustness and portability are further showcased. By open-sourcing all design files, preparation protocols, documentation and software, a useful, accessible, and self-contained neural interfacing solution is then delivered, catering to the expanding needs of both traditional and non-traditional *in vitro* applications.

## 2. Platform overview

Here, a system-level overview of our hardware and software is provided (Fig. 1). We start by describing the recording platform, its basic configurations and extensions. We then discuss its deployment in the lab and managing software. Finally, we compare our design with existing alternatives.

**Figure 1:**
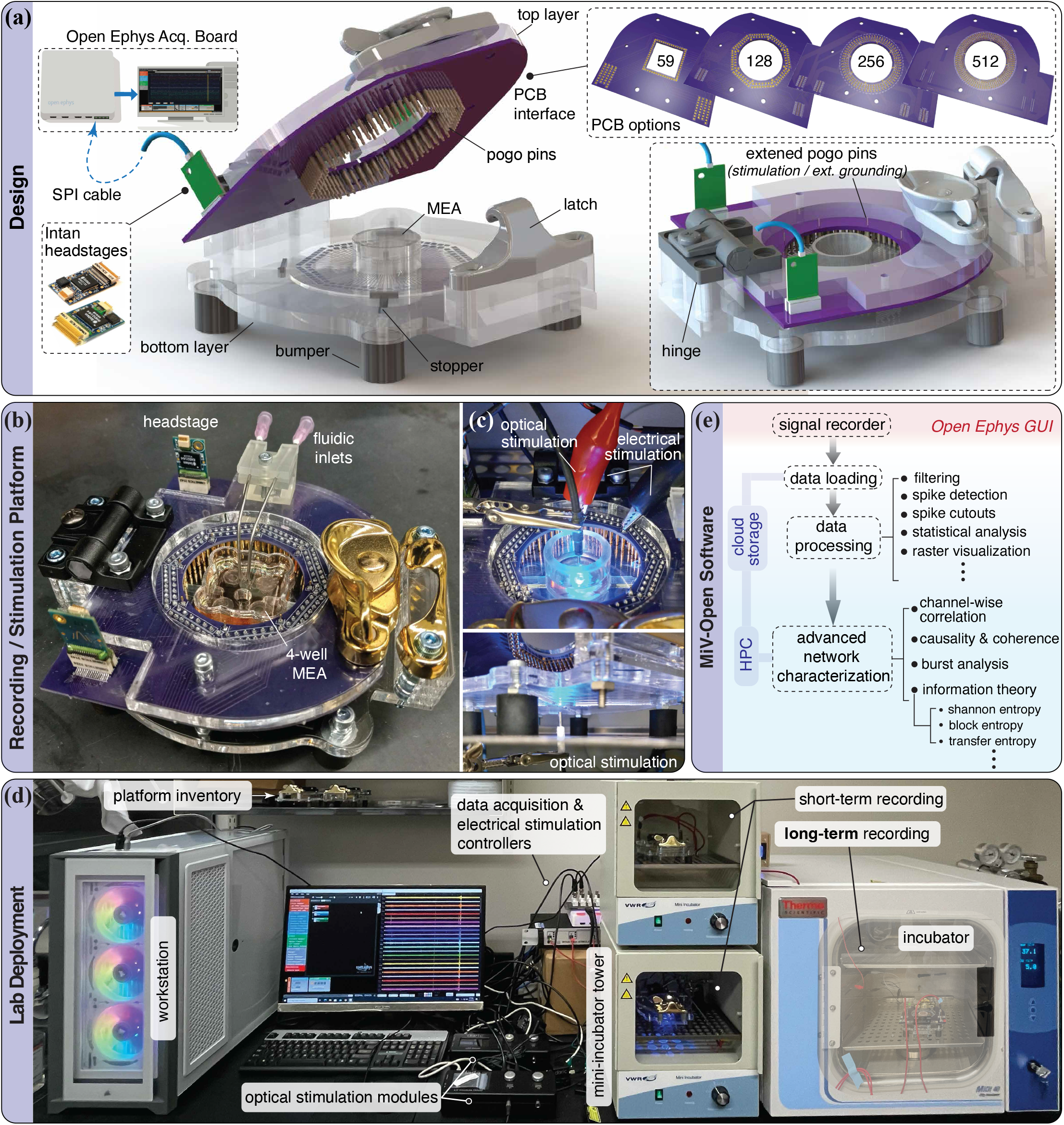
System overview. **(a)** CAD illustration of our recording platform, assembled from low-cost materials and commercially available parts. The platform is designed to be compatible with a series of PCB options across a variety of different recording capacities. The full list of components used to construct the hardware system can be found in the SI. **(b)** Assembled 128-channel recording platform. **(c)** Stimulation setup. Electrical stimulation can be performed by connecting a pulse generator (e.g. Stimjim) to extended pogo pins through a direct connection. Alternatively, for finer control, the Open Ephys terminal can be replaced with an Intan terminal, enabling bidirectional recording/stimulation across up to 128 channels. Optical stimulation is achieved by placing optical fibers on either side of the MEA. **(d)** Lab deployment of the recording system. Recordings are performed inside an incubator, while the data acquisition board and various external modules are set outside of the incubation environment. (e) Overview of the customized software package for data analysis, including an interface module for loading continuous or binary data recorded using the Open Ephys GUI, along with a variety of commonly utilized functions relevant for data processing and network characterization.

### Recording platform

Central to our system is a platform that hosts MEA-chips and PCB interfaces to form a complete signal chain, from neural substrates to data acquisition terminal and downstream processing (Fig. 1a). The platform consists of two layers of laser-cut acrylic board that are connected by a plastic hinge and can be locked in-place through a latch. This design eases loading and unloading of the chips and allows for rapid changes in recording configurations. The bottom layer also accommodates four sliding guides and 3D-printed stoppers, to conveniently center and hold chips of different sizes and shapes. Underneath the bottom layer, four soft rubber bumpers are affixed to insulate the platform against vibrations, minimizing recording noise. Secured to the top acrylic layer, a PCB provides the signal interface between the MEA chip and the Open Ephys data acquisition terminal. To ensure a firm, yet compliant contact with the MEA, arrays of spring-loaded contact (pogo) pins are soldered to the bottom of the PCB. Upon loading a chip, the top layer is closed and locked with the pins gently making contact with metal pads patterned on the perimeter of the MEA. These pads, and the electronic tracks emanating from them, are designed to connect one-to-one with micro-electrodes located at the center of the chip, where neural substrates are plated and incubated. Signals sampled at 30kHz at each electrode are then transferred through these tracks, pogo pins and PCB before being received and processed by amplifier headstages located on the rear edge of the PCB. Intan Technologies headstages, connected to Open Ephys terminals through serial peripheral interface (SPI) cables, are utilized here because of their wide compatibility.

#### Supported recording capacities

With versatility in mind, our system supports a range of recording capacities and configurations. The Open Ephys board allows the connection of up to 4 SPI cables, each capable of handling up to 128 digital data streams, setting the maximum capacity to 512 recording channels. This translates into high-density chips of 512 microelectrodes. Practically, in certain contexts, it may be desirable to employ fewer recording channels. Indeed, fewer channels imply fewer headstages (significantly reducing costs), simpler MEAs and PCB designs, as well as less computationally intensive data processing and analysis. Our system then provides built-in modularity to mix-and-match Intan headstages of variable capacity (32, 64, 128 channels), through a series of swappable PCB options supporting 59 (compatible with the widely used Multichannel system 60MEA series), 128, 256 and 512 recording channels. An example of 128-channel recording system, employing two 64-channel headstages, is illustrated in Fig. 1b. Assembly details for each standard are found in the SI.

#### Extension to electrical and optical stimulation

Our system supports both electrical and optical neural simulation to enable multimodal inputs to the neural tissue. Through extended pogo pins on the PCB (Fig. 1a), designated electrodes can be directly connected with an external electric source for stimulation. This allows the utilization of low-cost stimulators, such as Stimjim [36] (Fig. 1c). Alternatively, if a large number of stimulation channels is desired for fine spatial control, a bidirectional Intan controller can be connected to the platform’s headstages, enabling the simultaneous stimulation of up to 128 channels (details in SI). Another powerful, widely adopted modality of neural interaction is optical stimulation. At the cellular level, the expression of light sensitivity to specific wavelengths is achieved via channelrhodopsin transfection [37]. Here, we employ 465nm LED and lasers from Doric Lenses connected to optic fibers to locally stimulate transfected neural populations. To this end, our platform is designed to be top and bottom accessible, with openings on both acrylic layers, allowing optical fibers to be placed underneath or above the MEA-chip (Fig. 1c). This design allows to flexibly combine recording, and electrical/optical stimulation, either concurrently or serially, to achieve the desired input/output protocol.

### Deployment in the lab

The platform deployment is presented in Fig.1d, illustrating its integration with a workstation, multiple data acquisition/stimulation controllers, optical/electrical stimulators, and incubators of different dimensions and specifications, all fitting within the space of a standard benchtop. With the largest dimension below 150mm, the recording platform, upon sanitization, can be easily placed and operated across incubators. Apart from providing the necessary environment for maintaining neural cultures, incubators also serve as noise canceling Faraday cages that minimize interference, thanks to their conductive inner surfaces. Assembled from inexpensive materials, the recording platform can be conveniently duplicated to allow parallel recordings. Short-term experiments (<2 hours), during which culture media evaporation and pH change are not a major concern (at least in the case of dissociated cell cultures), are performed in a mini-incubator tower stack. Maintaining only temperature, these incubators are low-cost, portable, and have simple chambers that can be easily reconfigured and sanitized. Several demonstrations of Section 4 have been carried out in this environment. For long-term recordings (>24 hours) instead, more capable incubators for the control of temperature, pH, and humidity are necessary. The required high-humidity levels, in particular, pose a significant challenge for electronics, leading to rapid oxidation and signal degradation. This is one of the reasons why long-term *in vitro* electrophysiology remains uncommon, given the investment associated with commercial setups. In our case, the separation between data acquisition terminal and on-site signal amplification, combined with the use of inexpensive components, reduces the liability of operating our platform in incubator atmospheres for extensive periods of time, minimizing financial losses in the event of electronics failure. A 24-hour experiment with no appreciable hardware degradation is reported in Section 4.

### Software

Handling large streams of electrophysiology data, particularly when a high number of recording channels or multiple platforms are used in parallel, introduces challenges in compressing/archiving, transferring, and processing collected information. For reference, a single 24-hour experiment carried out with a 512-channel platform sampling at 30kHz produces approximately 4TB of data. In light of this rising demand, we have developed a cloud computing solution that simplifies the management of terabyte-size data and streamlines post processing. Our software offers a user-friendly interface for experimentalists who desire to construct processing pipelines and view results on their local desktop, while power-users can utilize the backend tools that support scalable high-performance computing (HPC) for custom, large-scale analysis. A software overview is presented in Fig. 1e.

The core of our software is a Pythonic pipelining platform specifically developed for maximizing flexibility and promoting consistency in experimental research. Our software provides a structured analysis template, while its backend incorporates essential features that are frequently used for analysis, eliminating the need for user implementation or maintenance. These features include caching, data pipelining, filtering, spike detection, principal component analysis (PCA), burst detection, criticality analysis, as well as an array of visualization functions. In addition, the software includes HPC support to integrate existing algorithms and pipelines into supercomputing clusters, enabling parallel processing and I/O capabilities to accelerate large-scale analysis. This template relies on common data structures and order of operation, ensuring compatibility among different data sources, methods, and algorithms. Furthermore, this approach modularizes the software, where users can selectively install only needed components as plug-ins, resulting in a compact and lightweight package.

To maximize impact, our software natively integrates with a variety of external packages of demonstrated utility, such as H5py with standard H5-data structure for scalable I/O [38], Aim UI for Slurm monitoring [39], Jupyter server for interactive GUI, PyInform/IDTxl for multi-variate analysis [40, 41], Globus APIs for synchronizing data stream, NeuralEnsemble [42, 43] or Kilosort [44, 45] for neurophysiology processing, and scikit-learn for machine learning [35]. Besides enabling the effective use of our platform, our software ecosystem aims at bridging the gap between experimental practice, data management and advanced analysis. While an in-depth characterization and demonstration of this software is beyond the scope of this paper, we note that all case studies presented here have been configured, analyzed and visualized using it.

### Comparison with alternatives and cost breakdown

With all major specifications illustrated, we summarize and compare our *in vitro* electrophysiology approach with other previously reported custom solutions. As illustrated in Fig. 2, from a recording system perspective our approach demonstrates the highest level of versatility, while being fully committed to open-sourcing both hardware and software. Further, and importantly, the robustness and utility of our system is comprehensively demonstrated across a range of neural substrates and stimulation settings, long-term and off-site recordings, as well as through extensions such as fluidic circulation and concurrent imaging.

**Figure 2:**
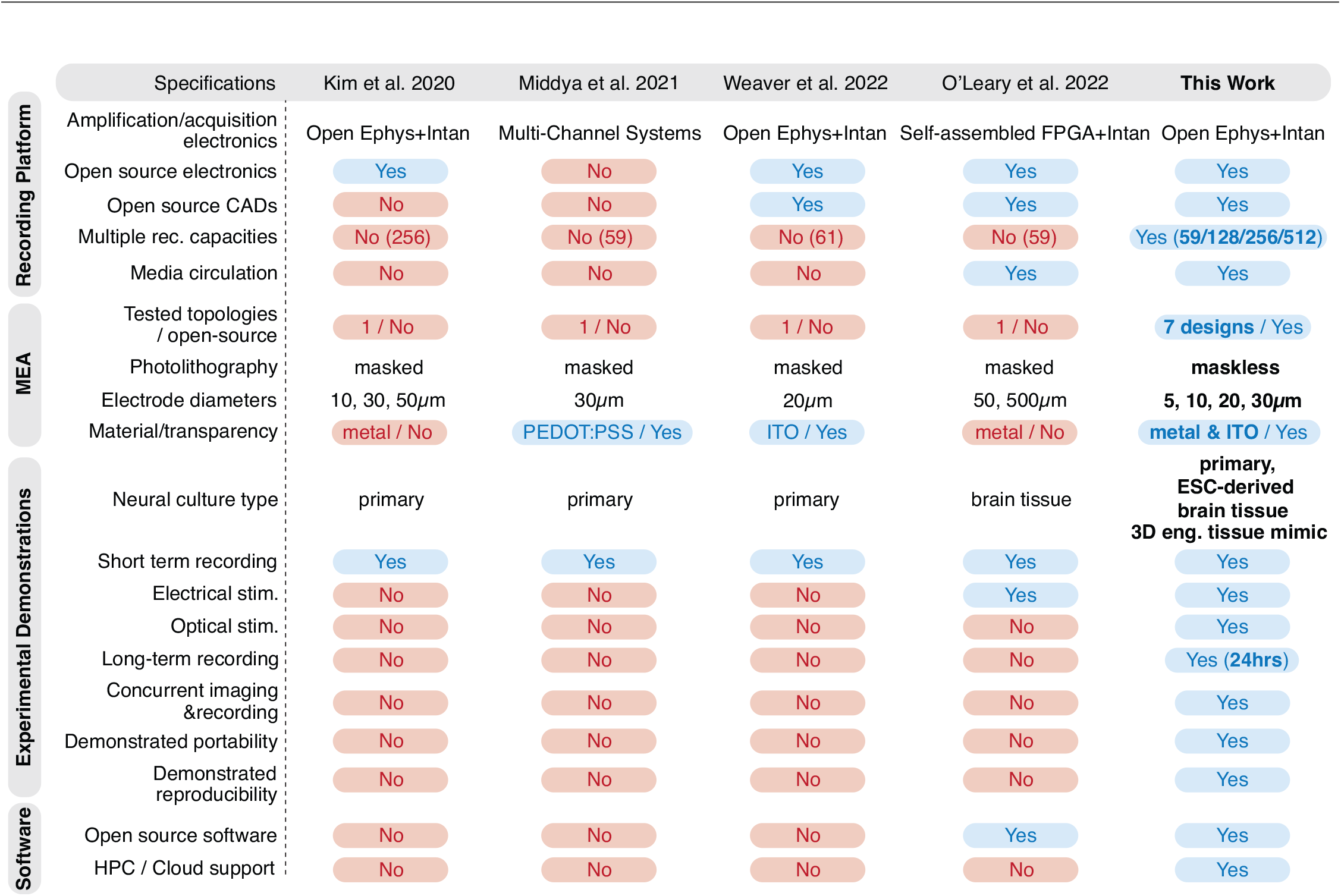
Comparison among recent custom electrophysiology approaches [27, 28, 29, 26]

A cost estimate is also provided. Incorporating open-source components as well as inexpensive, accessible materials, our platforms can be assembled at a fraction of the investment necessary for a comparable commercial product. Indeed, our platforms range from approximately ∼$2,500 for a 59-channel system to ∼$12,000 for a 512-channel device (both including the Open Ephys terminal), delivering ∼10-25X cost reductions, depending on vendor and comparative specifications. Given the modularity of our approach, this price advantage becomes even more significant considering that a single fabricated system can be reconfigured to satisfy a number of different or evolving needs. Details of the cost breakdown can be found in SI.

Finally, we emphasize that all CAD designs, manufacturing and assembly instructions, software and user documentation are made available with links to permanent repositories (Material and Methods and SI).

## 3. MEA-chip microfabrication, design and characterization

In keeping with an approach centered on versatility, here we focus on the realization of custom MEA-chips. We adopt a microfabrication process based on maskless photolithography. This technique enables the direct patterning of any 2D topology imported from CAD files, bypassing the need of dedicated photomasks for each design. Chips can then be customized without additional manufacturing steps, reducing fabrication time and cost. Further, this protocol allows significant flexibility in terms of constitutive materials, which we demonstrate and leverage to manufacture transparent MEAs for optical applications, as described in Section 4.3.

### Microfabrication

An overview of the adopted fabrication process is illustrated in Fig. 3a, while detailed information about tools and settings is provided in Material and Methods and SI. We start by spin-coating a thin layer of photoresist (PR) onto a clean borosilicate glass wafer. Glass is selected as substrate material because of its transparency, chemical stability, and strength. We employ the Heidelberg MLA 150 Maskless Aligner to pattern custom MEA designs on the PR layer via a direct-writing laser source. The processed wafer is then submerged in developer solution to dissolve the exposed PR and reveal the MEA topology. Next, constitutive materials of the MEA are uniformly deposited on the developed wafer through sputtering. Most of the MEAs employed in this study consist of two metal layers of titanium (Ti) and platinum (Pt), with a total thickness of 100nm. Here, Pt serves as the main constitutive material because of its good conductivity and relative ease to obtain and utilize, while the Ti layer is used for enhancing adhesion between Pt and wafer. An alternative process is also discussed in Section 4.3 for the deposition of thin films of indium tin oxide (ITO), which has been previously demonstrated in fully transparent MEAs [29]. Subsequent to the sputtering process, materials outside of the MEA pattern are lifted-off through sonication in acetone.

**Figure 3:**
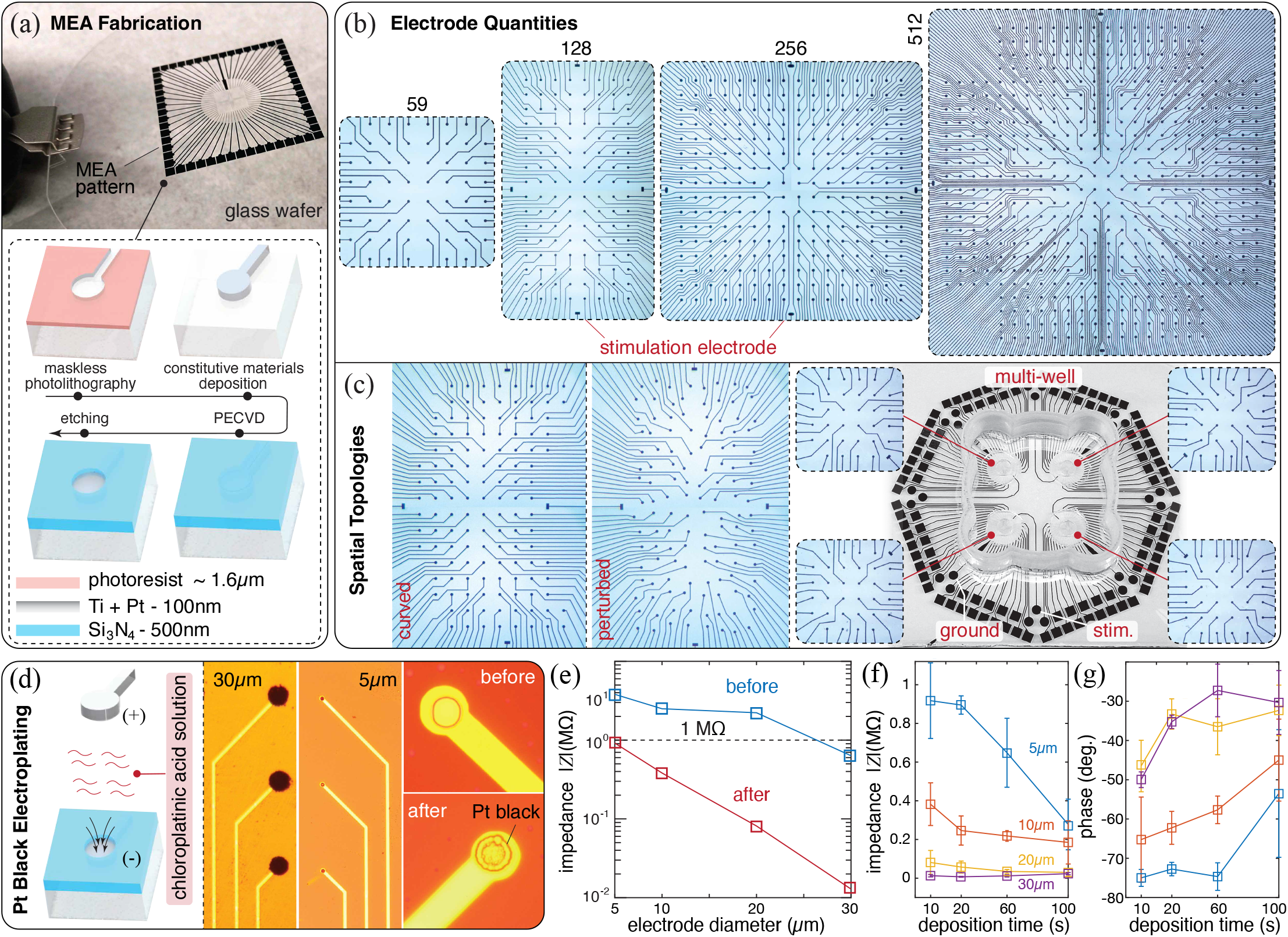
MEA-chips fabrication, design, and characterization. **(a)** Overview of the MEA fabrication process using cleanroom techniques. **(b)** Custom MEA designs of variable capacity (59, 128, 256 and 512 electrodes). Electrodes are 30*μ*m in diameter. The 59-electrode version replicates the design of the widely employed commercial Multichannel System 60MEA200/30 series. **(c)** Three 128-channel MEA designs with customized topologies. The curved layout presents a curvature of ∼0.35mm^*−*1^, matching the typical morphology of mouse cerebral cortex [46]. The perturbed layout entails random electrode displacements of up to 50*μ*m from the standard, rectangular layout of (b). **(d)** Platinum black deposition reduces electrodes impedance. Microscopic images are taken after deposition for both 30*μ*m and 5*μ*m electrodes. Close-ups illustrate the granular structure that Pt forms on the electrode. **(e)** Impedance magnitude before and after Pt deposition across electrodes of different sizes. **(f)** Time-deposition dependent impedance magnitude reduction. **(g)** Time-deposition dependent change in impedance phase.

Next, we apply the passivation layer that encapsulates the entire MEA, with the exception of the electrodes and contact pads. To do this, the wafer is first deposited with a 500nm thick layer of silicon nitride (Si_3_N_4_), through plasma-enhanced chemical vapor deposition (PECVD). Then, a second photolithography process is performed, creating a patterned PR layer on top of the Si_3_N_4_, with openings at the electrode and contact pad locations. The exposed Si_3_N_4_ is then removed by reactive-ion etching, using the PR layer as a mask. The PR layer is subsequently stripped using acetone. The overall result is a chip in which electrodes and contact pads are exposed for recording and stimulation while connecting wires are insulated. Finally, the wafer is diced into chips of desired shapes.

### Design

In Fig. 3b,c we demonstrate the versatility of the above approach through the realization of a variety of MEA designs. Figure 3b illustrates a series of 30*μ*m-electrode chips of increasing resolution spanning 59 to 512 channels arranged in regular rectangular patterns. Each of these MEAs presents a contact pad layout compatible with one of the PCB interfaces of Fig. 1a, synergistically enabling the rapid reconfiguration of the platform. Besides the number of electrodes, the chip layout can be modified as well. Figure 3c illustrates three designs, of which the first two have not been previously reported (either commercially or through customization): a curved MEA arrangement optimized to conform to the natural morphology of the mouse cortex [47], and a perturbed layout in which electrodes are randomly displaced around uniform grid locations, to reduce subsampling and aliasing in the analysis of neuronal avalanches [9], as theoretically predicted in [48]. The third example implements a multi-well topology, allowing to host and monitor four independent cultures at once for parallel testing. The use of several of these layouts is demonstrated in the Results section.

### Characterization

Prior to deployment for cell seeding, electrical properties of the chip array are characterized, and their impedance is reduced through platinum black deposition. This is a commonly employed electroplating technique for which platinum in chloroplatinic acid solution forms granular structures that adhere to the electrodes, upon current application [49, 50] (Fig. 3d, further details in Section 6.1). The process increases the electrodes’ effective surface area, lowering impedance and thus improving the capability of detecting neuronal signals.

In order to gain operational insight, we characterize the dependence between impedance and electrode size. We employ a specially designed chip (again leveraging the versatility of our fabrication approach) in which electrodes of four distinct diameters (5, 10, 20 and 30*μ*m) are patterned in separate quadrants (SI). Figure 3e shows that, while impedance is generally larger for smaller electrodes (as expected), in all cases the use of platinum deposition leads to reductions of at least one order of magnitude, yielding values below the neural signal detection threshold of *<* 1MΩ. This implies that all of these electrode sizes are viable, an additional asset relevant in specific applications. We also explore how impedance decreases relative to deposition time (Fig. 3f). Our results suggest that longer deposition time (of up to 100s) is needed for smaller electrodes (5-10*μ*m) to achieve the ideal value of ∼200kΩ [51]. Finally, we quantify the phase of the impedance (which is a complex variable) to confirm its negative value, that is to confirm the capacitive property of our electrodes. As can be seen in Fig. 3g, this is demonstrated across all cases, rendering our chips well-suited for electrophysiology.

## 4. Results

Through a variety of 2D and 3D neural systems derived from a broad number of sources, we illustrate the functionalities of our hardware and software both on-site and off-site, involving multi-modal stimulation, concurrent imaging and fluidic support. All demonstrations entail the use of a 128-channel platform configuration, selected because of its exact compatibility with both Open Ephys and Intan bidirectional modules. However, results relative to our other configurations can be found in the SI.

### 4.1 2D neural systems

We first test our interface via 2D cultures from both mouse embryonic stem cell-derived motor neurons (ESC-MNs) and dissociated mouse primary neurons (PNs). These cultures are prepared following different dissociation/differentiation protocols but share a conserved seeding procedure. Seeding is directly performed on MEAs that are surface treated to enhance cell attachment (protocols details in Section 6.2). Throughout, ESC-MNs represent our main cell line, as it endogenously expresses Channelrhodopsin-2 (ChR2) enabling optical stimulation [53], and because it can be differentiated into motor neurons that co-express enhanced green fluorescent protein (eGFP). This conveniently allows us to monitor culture conditions during preparation and maturation. Further, recorded data can be more easily interpreted with the aid of fluorescent microscopy. Results in this section are then shown for ESC-MNs, however, a parallel study with PN cultures (without GFP or optogenetic stimulation) is reported in the SI.

We seed ESC-MNs on two 128-channel MEAs, realized with either 30*μ*m or 5*μ*m electrode diameters (Fig. 4a), and begin recording spontaneous neural activity 7 days after seeding. Each recording consists of 2 minutes of data and is repeated every other day, at the same time, for 20 consecutive days. Raw neural signals are recorded using Open Ephys and are then processed offline through our software, for filtering and spike detection. The spike raster of Fig. 4b showcases spontaneous neural activity from the 30*μ*m MEA, where single unit activity is plotted across channels. The use of high-density MEAs helps reveal intrinsic network properties, reflected here in highly synchronized population bursts, which would not be detectable at the single site level (e.g., patch clamp electrophysiology). Waveforms from all recording channels are sorted using PCA to isolate signal fingerprints matching characteristic neuron spike shapes (cutouts of Fig. 4b). The performance stability of our system is illustrated by plotting daily average spiking rates across all channels over 4 weeks, where a typical *in vitro* neuron maturation-degradation curve is observed. As can be seen in Fig. 4c, for MEAs of both electrode sizes, average spiking rates first increase during network maturation, to then plateau and gradually decrease as cells begin to age. We also quantify the quality of our recording setup by determining the mean signal to noise ratio (SNR) across channels. As seen in Fig. 4d, spiking channels (95% of channels for 30*μ*m MEA and 70% for 5*μ*m MEA) consistently exhibit an SNR*>*5 after filtering, on-par with reported performance of commercially available devices [54].

**Figure 4:**
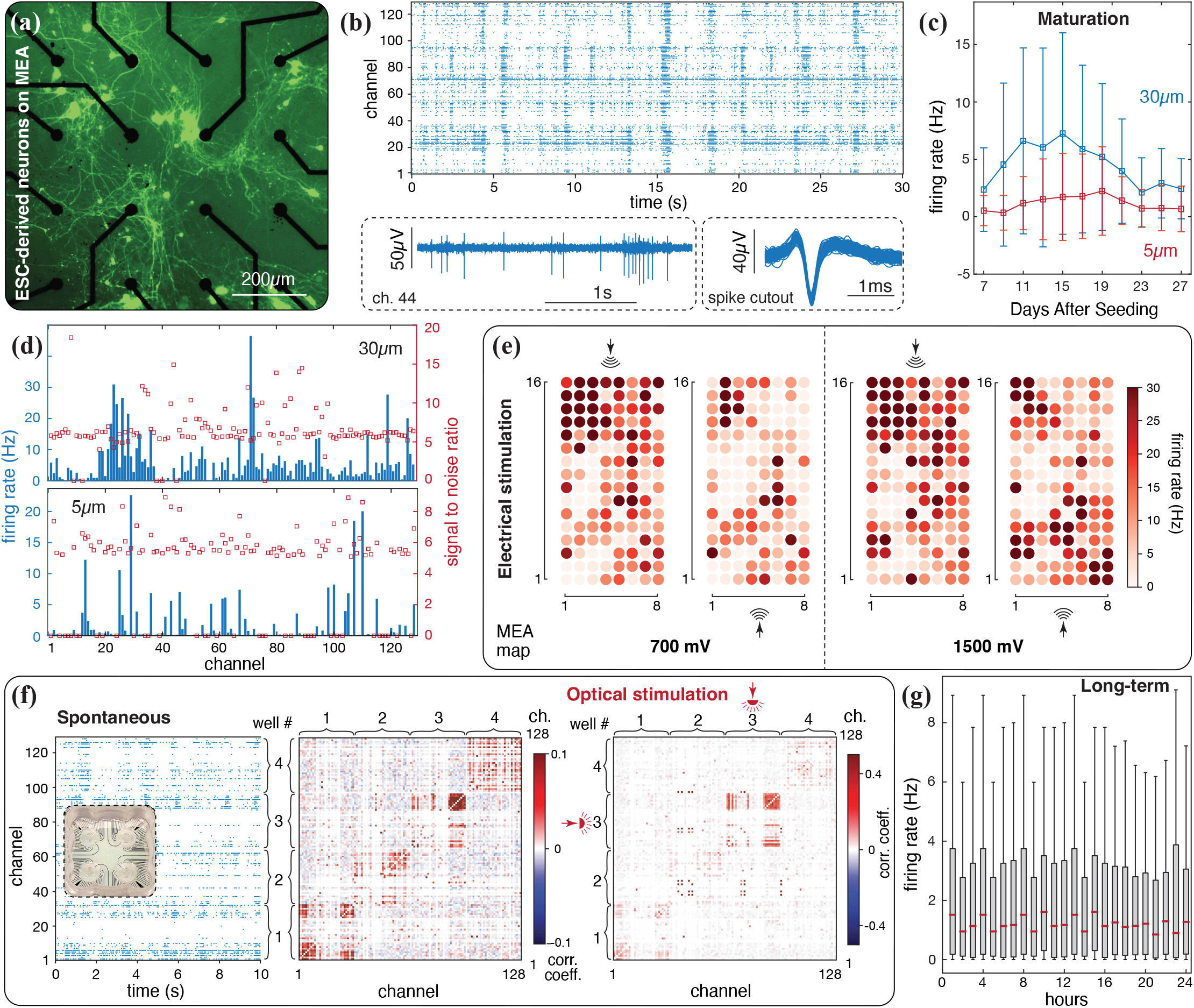
2D neural systems. **(a)** Fluorescence microscopic image of eGFP ESC-MNs seeded on MEA. **(b)** Raster plot shows the spontaneous neural activity recorded from a rectangular MEA layout (Fig. 3b) of 30*μ*m-electrode on day 17 after seeding. Three seconds of filtered data from a representative channel reveal individual spikes as well as a burst event. Filtering is performed using a third-order bandpass Butterworth filter with cutting frequencies at 200 and 3000Hz. Spikes are overlaid after sorting. **(c)** Averaged firing rates across all channels over 2 minutes, as a function of days after seeding. Results from MEA chips of 30 and 5*μ*m electrodes are reported. Error bars indicate standard deviation across channels. **(d)** Averaged firing rate of each recording channel on day 17. The dataset is used to calculate the signal to noise ratio (SNR), defined as the ratio between the mean spike peak amplitude and the standard deviation of the full signal. **(e)** Spatial activity propagation in response to localized electrical stimulation, at two different stimulation sites (top and bottom) and with two different intensities (700mV, 1500mV). Each stimulation experiment entails 6 to 8 trains of 20Hz biphasic stimulation pulses, with each train lasting for 1 second. Biphasic pulses present a positive-to-negative transition, where each phase has a duration of 400*μ*s. These parameters are selected based on [52]. Data are recorded from the 30*μ*m MEA of Fig. 3b, on day 17 for the 700mV tests and on day 19 for the 1500mV test. **(f)** ESC-MNs cultured on a 4-well MEA. Spontaneous activity is recorded (left raster plot) and channel-wise correlation coefficients are computed to demonstrate separation between different clusters. We then proceed with selective optical stimulation of well #3. Stimulation lasts 1min with a 1s pulse train delivered every 5s. Each train is comprised of pulses at 40Hz, 20% duty cycle and 1A current amplitude. The experiment is performed on day 13 using a 465nm LED source acquired from Doric Lenses. **(g)** Long-term recording for 24 hours. Box plot shows the spontaneous activity recorded for the first 12mins of every hour. The red dash denotes the median value, the box indicates the range from the first quartile (*Q*_1_) to the third quartile (*Q*_3_), and the vertical line extends from the box by 1.5 times of the inter-quartile range (IQR = *Q*_3_ − *Q*_1_). Outliers beyond the range of the vertical line are not plotted.

On top of spontaneous activity recording, we consider simultaneous multi-modal stimulation. While this is a useful paradigm in neuroscience for studying synaptic potentiation [55], it is also a pre-requisite to encode inputs for potential computing applications. As an example of electrical stimulation, we consider a single-well MEA seeded with ESC-MNs, and apply brief, biphasic electric pulses (Stimjim [36]) at a designated location. The heat-maps of Fig. 4e show the effect of such stimulations, visualizing the average neural firing recorded by each microelectrode. We find that within our cultures, firing rates of neurons surrounding the stimulation site are enhanced, with levels of activation proportional to the intensity of the stimulus.

We proceed by pairing optical stimulation with a 4-well chip design, to illustrate a parallel environment for control and selective stimulation experiments. To this end, ESC-MNs are seeded into four independent clusters on the MEA of Fig. 4f. Spontaneous activity is first recorded and processed to compute network correlations within wells and across wells. We see that recordings within the same cluster show high correlation scores. We then selectively apply optical stimulation to one well (#3), obtaining a highly synchronized response compared to the other wells.

Finally, we consider long-term electrophysiology applications. Complementary to short-term recordings that reveal transient and fast neural responses, longer recordings are necessary for investigating prolonged and slow plastic behaviors, such as facilitation, habituation, and long-term potentiation. However, as underscored in Section 2, performing long-term recordings inside a high-humidity environment challenges the recording system with accelerated electronic degradation. We reduce liability via a modular design for which most of the delicate electronics are kept outside the incubation chamber. It should further be noticed that in case of failure, financial losses are minimized given our system’s low-cost. We then demonstrate the continuous monitoring of ESC-MNs cultures over 24 hours. As illustrated in Fig. 4g, an overall consistent level of spontaneous neural activity is observed, showing no sign of either hardware or culture-wise degradation throughout the recording. We also note that prolonged recordings inevitably produce high volumes of data (in the TB range), rendering post-processing via standard PC workstations cumbersome and time-consuming. This motivates the extension of our software to include functionalities for streamlining large-scale post analysis through cloud storage and HPC. Results in Fig. 4g are obtained by deploying such functionalities on the supercomputing facility Frontera at the Texas Advanced Computing Center.

### 4.2 Higher dimensional neural systems

While 2D cultures allow for initial characterization and understanding of a networked cellular system, their lack of 3D organization does not fully capitalize on neurons’ potential for enhanced connectivity, compute density, or miniaturization, nor can they recapitulate physiological architectures. Here, we demonstrate our platform in a higher-dimensional context, by considering both *ex vivo* tissue and 3D engineered mimics.

#### Organotypic brain slices

Organotypic cultures are prepared from slices of rat cortex that are 450*μ*m thick. While these compress to about 100*μ*m during incubation, they nevertheless preserve an intrinsic connectivity structure which is lacking in the 2D monolayer cultures described above. At the same time, brain slices do not possess a clear 3D shape like the tissue mimics we will discuss below. To highlight this distinction, we consider organotypic cultures to be of intermediate dimension (2.5D). Relative to 2D cultures, experimentation with higher dimensional systems poses additional challenges such as ensuring that the tissue is broadly and firmly in contact with the microelectrodes, as well as provisions for continuous media, oxygen or drugs replacement. To address these challenges, we augment our system through a 3D printed fluidic interface that is modularly integrated via accessory posts protruding from the top acrylic layer (Fig. 5a). The printed apparatus hosts a pair of L-shape fluid delivery cannulas, connected to a peristaltic pump for continuous media delivery and aspiration. This can be further extended to support a ‘plug’ system whereby a biocompatible nylon filter mesh can be gradually lowered towards the tissue to apply a gentle pressure onto it, thus enhancing contact across the electrodes surface.

**Figure 5:**
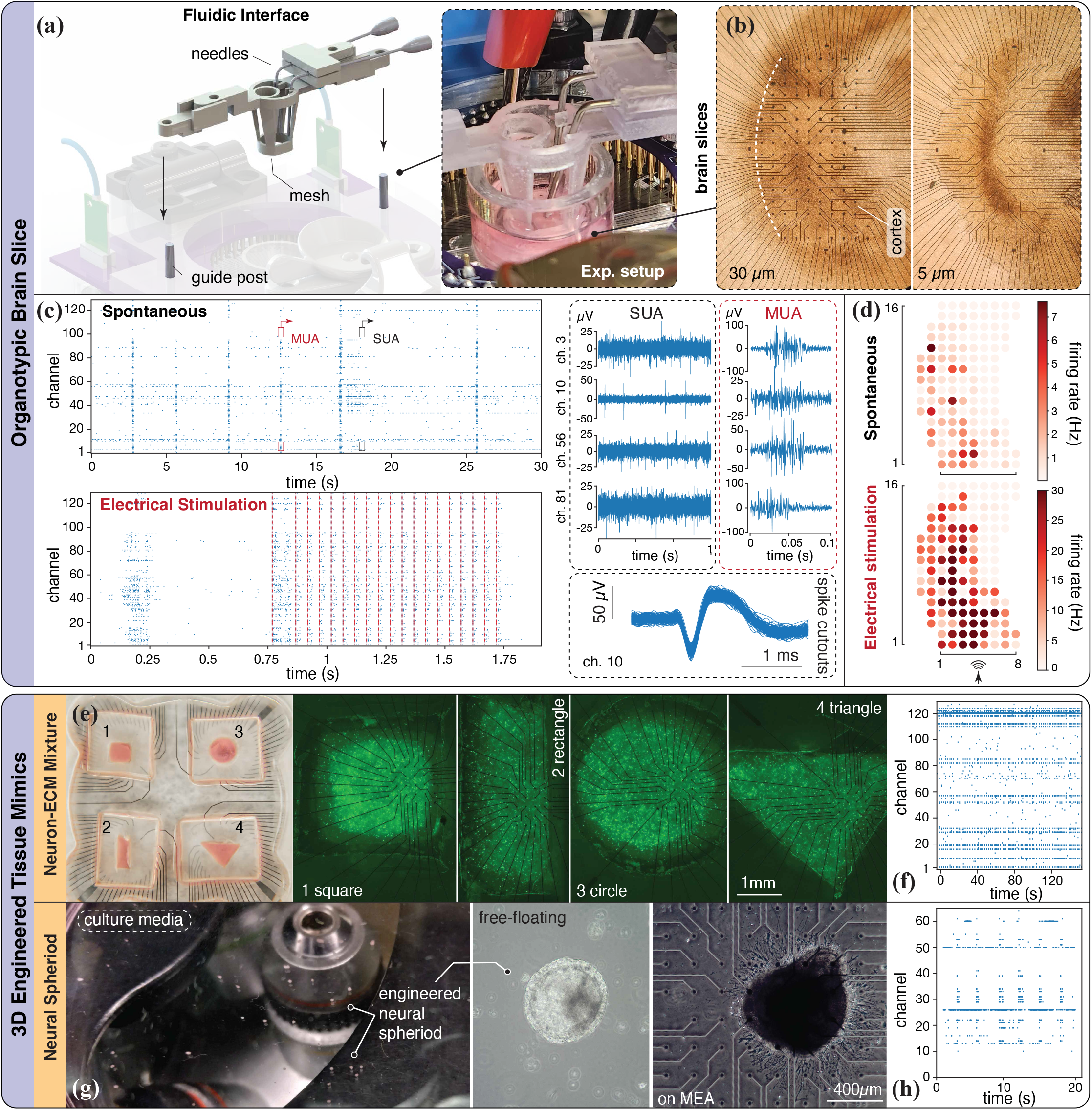
Higher dimensional neural systems. **(a)** Platform integration with an add-on fluidic interface module for brain slice recordings. The add-on module is made of bio-compatible stereolithography material using a Formlabs 3D printer. Before the experiment, the printed parts are autoclaved and sanitized to ensure nontoxicity to the brain tissue. Hypodermic needles used for fluid flow are grounded to the recording system to minimize noise levels. **(b)** Microscopic images of organotypic brain slices placed on curved MEAs with either 30 or 5*μ*m electrode diameters. Electrodes are arranged to match the curvature of the cortex region, which is the shaded area around the center of the slice. **(c)** Spike raster of spontaneous tissue activity, measured around 10mins after transferring onto the MEA. A raster plot corresponding to electrically stimulated activity, measured around 15mins after transferring, is also reported. Each red dashed line indicates a single biphasic pulse (applied at the bottom stimulation site). Snapshots of filtered neuronal signals during spontaneous recording showcasing single-unit and multi-unit activities. Cutouts are plotted to demonstrate the detection of characteristic spike shapes. **(d)** Firing heatmaps of spontaneous and stimulated activities. **(e)** 3D engineered neural tissue mimics (NTMs) made of cell-ECM mixture. Custom multi-well MEA hosts PDMS molds of 4 different shapes, demonstrated in various controlled NTMs geometries. GFP signals illustrate motor neurons distribution within each tissue. **(f)** Spontaneous activity recorded on day 7, from an NTM sample seeded on a 128-electrode, rectangular MEA. **(g)** Engineered neural spheroids. Microscopic images are taken to visualize free-floating spheroids and after transferring onto a commercial MEA (used here to confirm cross compatibility with our platform). **(h)** Spontaneous activity recorded from the same sample.

Using this particular accessory, we perform longitudinal recordings in mature organotypic cortex cultures. As previously discussed, this preparation is commonly employed because it preserves much of the intrinsic tissue structure found *in vivo*, like cortical layers and corresponding cell types. Further, it facilitates in-depth culture manipulation via localized drug delivery, optical control and cell identity imaging, which are difficult to perform *in vivo*. Cortical tissue is dissected from rat pups on postnatal day 5. These slice cultures are incubated for 26 days before transferring the slices to the MEA-chips for recording (protocol in Section 6.2). For this application we utilize the curved MEA of Section 3 to conform to the natural morphology of the region of interest (Fig. 5b). Neural activity recorded from a 5*μ*m-MEA sample is presented in Fig. 5c, illustrating spontaneous spiking events as a raster plot, together with corresponding waveforms (spike cutouts). We use our software to isolate both multi-unit activity (MUA) and single-unit activity (SUA) across the electrode array, and find that MUA is highly synchronized across the cortical culture, consistent with significantly correlated bursting activity.

In addition to spontaneous recordings, we employ the same biphasic stimulation protocol of Section 4.1 to study the tissue neural response. Activity in the sample is found to be strongly driven by the electrical stimulation (the red dashed line overlaid to the raster plot) across the entire 128-channel array. We note here that the use of curved MEAs also allows for consistent alignment of electrode sites to anatomical points of reference across the tissue. This enhances yield and increases usability, making it easier to reliably place and align the tissue to enhance experimental consistency across samples. Good alignment and yield are demonstrated in Fig. 5d, where recorded activity (firing rates) is mapped to the electrodes’ physical locations.

#### Engineered tissue mimics

While organotypic slices are representative of the intrinsic connectivity of brain tissue, engineered mimics potentially allow the realization of 3D neural architectures of desired size, topology and composition. This, combined with custom MEAs for interfacing, provides a unique opportunity to extend applications from neuroscience to engineering devices for sensing, processing and computing. We present here two different methods of bio-fabricating 3D engineered neural tissue mimics (NTMs), and demonstrate their integration in our platforms.

We first extend our 2D monolayer cultures to 3D NTMs by mixing ESC-MNs with 6mg/ml Matrigel (ECM). Here, we showcase the control over our NTM geometry by seeding the cell-ECM mixture into PDMS (polydimethylsiloxane) molds placed on our multi-well MEAs (Fig. 3c). Molds are fabricated with cavities of different shapes, allowing the mixtures to polymerize into 3D neural constructs of prescribed configuration (Fig. 5e). To confirm the success of the NTM fabrication, fluorescence imaging is performed to visualize live neurons’ GFP signals. As illustrated in Fig. 5e, neurons are evenly distributed within the constructs, and network formation is observed. We then deploy our platform for electrophysiology recording. Figure 5f illustrates the activity obtained from an NTM sample, where spikes and synchronized events are detected across multiple channels. We note that fewer active channels are seen here compared to 2D neural cultures. Indeed, since neurons are distributed in three-dimensional space the number of cells residing at the bottom of the tissue is reduced, rendering activity less detectable by the electrodes.

Alternatively, NTMs can also be realized through the creation of 3D neural spheroids. Formed from embryonic stem cells, and further differentiated into neural lineage, these spheroids recapitulate physiologically relevant features such as cellular and extracellular matrix composition [56]. We fabricate the neural spheroids by first obtaining embryoid bodies following the standard SFEBq method [57], and then differentiating them into cortical lineage by selective Shh (via cyclopamine) and Wnt pathway (via IWP-2) inhibition. The resulting spheroids are maintained in a suspension culture for 20 days before seeding on MEAs for recording (Fig. 5g). Here, in order to showcase the native platform compatibility with commercial MEAs, we specifically employ one from the Multichannel system 60MEA series. Figure 5g depicts a microscopic image taken on day 7 after seeding, illustrating a spheroid extending neurites on the commercial MEA. Corresponding recordings on day 12 after seeding demonstrate the successful detection of neural firing (Fig. 5h).

### 4.3 Concurrent calcium imaging

A unique aspect of a fully-customizable system is that its components can be continuously adapted to comply with existing industrial standards, or new standards as they emerge, while retaining the same core infrastructure. We showcase this here by reshaping our platform to allow concurrent calcium imaging. We alter the design of Fig. 1a to match the dimensions of a standard 96-well microplate (Society for Biomolecular Screening, SBS), enabling the integration with inverted microscopic chambers. As seen in Fig. 6a, a new PCB interface that accommodates a single 128-channel recording headstage (rather than two 64-channel headstages as in Fig. 1b) is mounted onto reshaped acrylic layers to deliver a compact, SBS-compatible layout.

**Figure 6:**
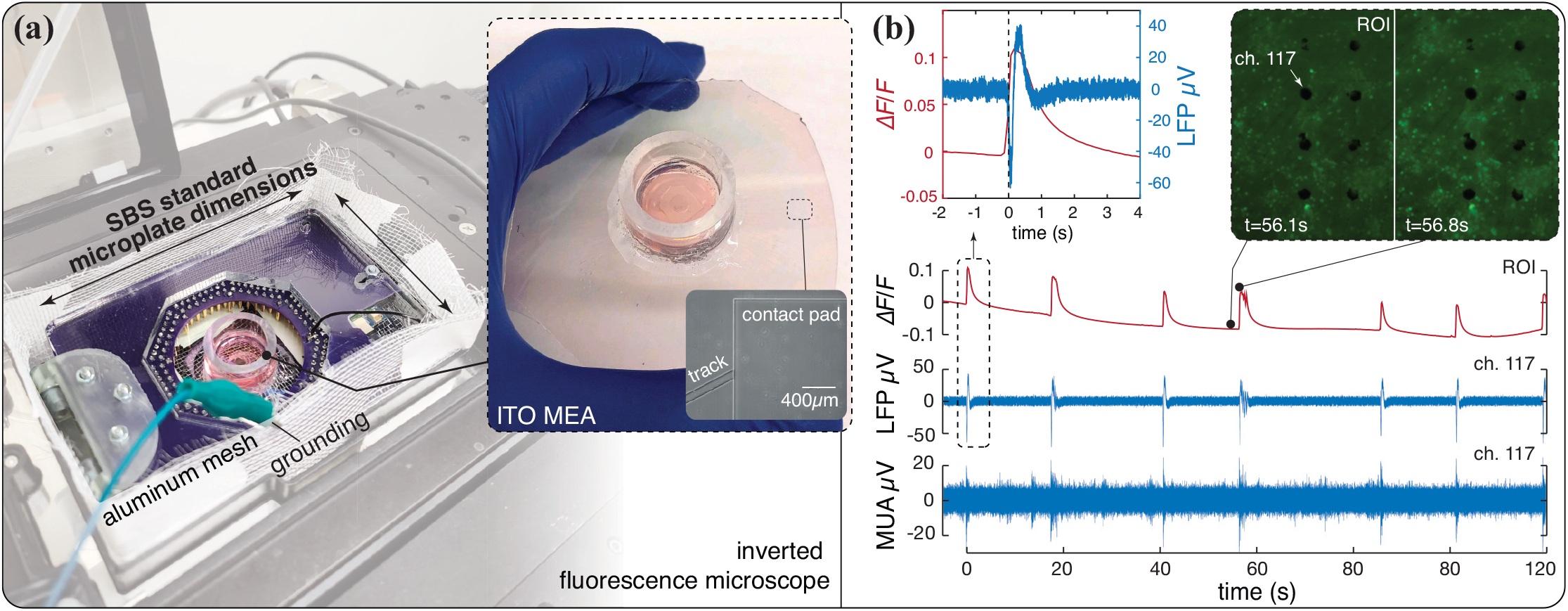
Alteration of the recording platform to facilitate concurrent electrophysiology and microscopic imaging. **(a)** Placing of the recording device inside an inverted microscopic chamber. A thin layer of aluminum mesh is fabricated to wrap around the device and serve as Faraday cage. ITO MEAs and PNs are utilized in this experiment. **(b)** Examples of simultaneously acquired fluorescence signal and electrophysiology data. Video analysis is performed within a ROI of 0.57 ×0.82mm, while electrical data are recorded from a channel located near the center of the ROI. Left inset: A peak in the ∆F/F_0_ signal and a LFP event share similar temporal characteristics, where a rapid initiation period (*<*150ms) is followed by a gradual restoring phase (1s for LFP and 3s for ∆F/F_0_). We align the two signals according to their first edge. Right inset: snapshots of the fluorescence signal before and during a burst event.

Upon integration within the microscope chamber (Fig. 6a), we proceed with testing the simultaneous electrophysiology and calcium imaging in 2D cultures of primary hippocampal neurons (PNs). PN cells utilized in this experiment are collected from embryonic rat brains (E18-E19) and cultured on MEAs following the same ESC-MNs seeding protocol. PNs are chosen because, unlike ESC-MNs, they do not express constitutive fluorescent reporters, and therefore allow the undisturbed visualization of the calcium signal. For this specific application we employ transparent MEAs. These are microfabricated following the same procedure described in Section 3, except for the addition of an annealing process after the ITO deposition to enhance transmittance and conductivity [58]. Annealing is carried out by placing the MEA sample in a vacuum chamber at 450°C for 1 hour, leading to ITO transmittance of ∼ 80% and resistivity of ∼ 4.5e^−4^Ωcm (see SI), enabling both optical electrical measurements.

We perform concurrent measurements 12 days after seeding. Cells are loaded with the calcium indicator Oregon Green 488, following manufacturer’s recommendations (Section 6.3). The loaded sample is then placed in our platform within the microscopic chamber, ready for measurement. After setting the laser source, video and electrophysiology recording of the culture’s spontaneous activity are simultaneously acquired, and offline analyzed. A representative example of post-processed signals is illustrated in Fig. 6b. Calcium activity is characterized through changes in fluorescent intensity (∆F/F_0_, defined in Section 6.3) within a region of interest (ROI), which in turn determines the microelectrodes to be considered. Each peak in the fluorescence plot corresponds to a culture-wide burst event, during which we can observe a higher-intensity calcium signal across the network, relative to the rest state (illustrated in the inset images). Detected electrical signals are processed separately to reveal local field potentials (LFPs) and MUA during bursts, demonstrating that recording is not affected by laser-induced artifacts. Further, LFPs and MUA are found to precisely align temporally, confirming simultaneous neural responses across modalities. In a context where genetic optical markers continue to provide new avenues to probe cellular dynamics at high-spatial resolutions, complementary high-temporal resolution electrophysiology is seen as a powerful feature.

### 4.4 Portability, robustness and reproducibility

Finally, we expand our discussion to emphasize our platform’s portability, robustness and reproducibility, key factors for broad dissemination.

Electrophysiology measurements are delicate and sensitive to hardware setup and testing environment, rendering the portability of recording solutions rarely reported or discussed (Fig. 2). Here, we demonstrate this capability through off-site recordings of organotypic brain slices prepared and cultured in Prof. Beggs lab at Indiana University Bloomington (presented in Section 4.2). To this end, a platform in use at the University of Illinois at Urbana-Champaign (UIUC) is disassembled and ground transported to Bloomington (∼180 miles, ∼3-hours drive). As presented in Fig. S5 in the SI, our system, after transportation, could be easily reassembled and deployed for experiments within half an hour. This demonstration compounds the above described case studies, further underscoring the robustness, versatility and accuracy of our systems across a variety of conditions.

These favorable features led us to collect nearly 1000 hours of recording experiments in one year, amounting to tens of TB of data. This ability has motivated external researchers to adopt our systems as an alternative to commercial devices, providing us with the opportunity of testing the independent reproducibility and implementation of our platforms. Following our open-source designs and protocols, a graduate student from Prof. Saif lab at UIUC, with little experience in electronics and fabrication and without supervision, was able to assemble a working system within two weeks (Fig. S7 in SI).

## 5. Conclusion

We have presented a versatile, scalable and multi-modal electrophysiology solution, delivering a customizable signal pipeline stretching from *in vitro* neural substrates to cloud computing. Our approach significantly lowers barriers to entry, through the open-sourcing of designs, software and protocols, and via (over) ten-fold cost reductions. The utility of our platforms is showcased through a comprehensive set of demonstrations involving a variety of cell types and stimulation modalities, two- and three-dimensional systems, concurrent imaging and long-term recording. This study provides a launching pad to broaden the electrophysiology domain, both in terms of users and applications, and motivates us to further innovate across scale and functionalities, paving the way to new biophysical discovery and *in vitro* technologies.

## 6. Material and Methods

### 6.1 MEA fabrication and Pt deposition

We fabricate our MEAs on a set of borosilicate glass wafers (Borofloat 33) ranging from 3 to 6 inches in diameter. The process starts with spin-coating of AZ5214E photoresist (PR) using a Headway PWM32 spin coater at a steady speed of 3000 rpm. Photolithography is then performed using the Heidelberg MLA 150 Maskless Aligner through a 375nm laser source with a dose of 210mJ/cm^2^. Developing is subsequently performed for 30s using 1:4 diluted AZ400K developer, followed by a 50s post bake at 120°C. Next, the AJA sputter coater series are leveraged for constitutive material deposition. Ti and Pt are both deposited at 3mT pressure, with respective power of 200W and 50W, while ITO is deposited at 5mT pressure, 80W power, and under the airflow of argon and 3% oxygen. Samples are then submerged in acetone for lift-off. We then employ Oxford PECVD to apply a 500nm passivation layer of Si_3_N_4_, which is produced with the supply of 20sccm of SiH_4_ and NH_3_ respectively, under 650mT pressure and 20W power. The second photolithography process is carried out using the same aforementioned parameter settings. The exposed passivation material is removed using Oxford Freon RIE with 30sccm CF_4_ as the etching gas. Finally, the sample is cleaned with acetone and diced using a wafer cutter. An illustration of our full fabrication process can be found in SI.

To electroplate the Pt black, we first prepare and mix 100mL of chloroplatinic acid that contains 1g of Hydrogen Hexachloroplatinic Hydrate, 0.01g of Lead(II) Acetate Trihydrate and 0.25mL of Hydrochloric Acid (Sigma-Aldrich) [49]. We add 1mL of mixed solution to each MEA and then electroplate utilizing a Keithley 6221 current source to supply a DC current of density 4 nA/*μ*m^2^, with the ground electrode being anode and all other electrodes being cathode.

### 6.2 Neuronal culture preparation

The functionality and versatility of our system and MEA are demonstrated through the recordings from a range of *in-vitro* neuronal cultures. This subsection presents the materials and protocols used to prepare each type of culture.

#### Embryonic stem cell-derived motor neurons

For the preparation of ESC-MNs, we culture optogenetic mouse ESC cell line ChR2^H134R^-HBG3 Hb9-GFP following an established protocol [59]. Briefly, mESCs are first cultured on a feeder layer of CF-1 mouse embryonic fibroblasts (Gibco), then the mESCs are suspended in an induction medium (advanced DMEM/F-12 and Neurobasal media with 10% KnockOut serum replacement, 1% L-glutamine and 1% penicillin-streptomycin) in a low-adhesion cell culture dish for two days to allow for the spontaneous formation of embryoid bodies (EBs). The EBs are then suspended in a differentiation medium (induction medium supplemented with 2% B-27, 1% N-2, 1% insulin transferrin selenium, 1*μ*M retinoic acid and 1*μ*M smoothened agonist) for up to 5 days. GFP expression is monitored daily to confirm the differentiation into motor neurons. After differentiation, the EBs are dissociated using Accutase and the resulting single-cell suspension is seeded on the MEAs at a density of 5000/mm^2^, following the seeding protocol detailed below. The neurons are kept in maintenance media (Neurobasal plus media with 2% B-27 plus, 1% L-glutamine, 1% penicillin-streptomycin). In the first 4 days after seeding, maintenance medium is supplemented with growth factors (BDNF, GDNF, CNTF, NT-3, Forskolin, IBMX) to promote neurite outgrowth and cell viability [60].

#### Primary neurons

Primary hippocampal neurons are dissected from time-pregnant rats at E18-E19, and put in MilliQ water (pH 7.4, 4°C) supplemented with 1.16% Na_2_SO_4_, 0.52% K_2_SO_4_, 0.24% HEPES, 0.18% D-glucose and 0.1% MgCl_2_ 6H_2_O. Hippocampal neurons are further dissociated and kept in MEM Eagle’s with Earles’s BSS w/o L-glutamine supplemented with glucose, 100mM sodium pyruvate, 200mM L-glutamine, and 100U/mL penicillin and streptomycin. Seeding and subsequent culture maintenance are similarly performed as for ESC-MNs.

#### Seeding on MEAs

To prepare the MEAs for culturing dissociated cultures (both ESC-MNs and PNs), MEAs are first surface-treated using oxygen plasma, followed by an overnight coating of 0.1mg/mL Poly-D-Lysin (PDL) (Gibco) at room temperature. After the PDL solution is removed, the MEAs are washed with phosphate buffered saline (PBS) and allowed to air dry completely. MEAs are then incubated with 20*μ*g/mL laminin solution (Sigma Aldrich) at room temperature overnight, and the coating solution is removed right before cell seeding without air-drying.

#### Organotypic brain slices

Following an established protocol[61], brains from postnatal day 5 rats are sliced into 400*μ*m slices containing somatosensory cortex. Slices are then incubated in a humidified atmosphere with 5% CO_2_ at 37°C in the culture medium, which consists of 2:4 Minimum Essential Medium, 1:4 Hank’s Balanced Salt Solution, 1:4 Horse Serum, 4mM glutamine and 1:100 penicillin/streptomycin. Half of the medium is replaced every 3 days. Two weeks after slice preparation, slices are removed from the incubator and placed on the curved MEA for recording. During recording, oxygenated culture medium is pumped into and out of the MEA well via a peristaltic pump (see SI). An inline heater is placed immediately before the well to warm the medium to 37°C. Both prior to and after each experiment, distilled water and isopropyl alcohol are run through the entire fluid line to clean them of any biological debris.

### 6.3 Calcium imaging

#### Loading calcium indicator

Calcium imaging is performed by first loading the neuronal culture with a cell permeable fluorescent calcium indicator, Oregon Green 488 BAPTA-1 (AAT Bioquest). To do this, the calcium indicator is diluted in the neural maintenance medium at a final concentration of 4*μ*M, and then supplied to the culture to replace the old medium. Subsequently, the culture is incubated for 40mins in a humidified environment with 5% CO_2_ at 37°C. Right before the measurement, the culture is washed twice with PBS and supplied with fresh medium.

#### Fluorescence intensity measurement

Spontaneous calcium activity is acquired by employing a fluorescent microscope (Nikon Eclipse Ti) and a fluorescein isothiocyanate (FITC) filter with excitation/emission wavelength of 490/525nm. Recording is conducted under a 488nm laser illumination and at an original frame rate of 100fps. We then down-sample the recorded video to measure the fluorescence intensity using the open-source software ImageJ. The variation of the fluorescence intensity is defined through the normalized form as ∆F/F_0_=(F-F_0_)/F_0_, where F and F_0_ are the instantaneous and initial fluorescence intensity, respectively.

## Supporting Information

Supporting Information is available from the authors.

## Acknowledgements

This study is jointly funded by National Science Foundation (NSF) EFRI C3 SoRo #1830881 (M.G.) and NSF Expedition ‘Mind in Vitro’ award #IIS–2123781 (M.G., J.B., H.K., T.S.). We also thank the Frontera computing project at the Texas Advanced Computing Center. Frontera is made possible by National Science Foundation award OAC-1818253.

